# ANTENNA, a Multi-Rank, Multi-Layered Recommender System for Inferring Reliable Drug-Gene-Disease Associations: Repurposing Diazoxide as a Targeted Anti-Cancer Therapy

**DOI:** 10.1101/192385

**Authors:** Annie Wang, Hansaim Lim, Shu-Yuan Cheng, Lei Xie

**Author notes:** Contributed equally to this work (Annie Wang, Hansaim Lim).

## Abstract

Existing^1^drug discovery process follows a reductionist model of “one-drug-one-gene-one-disease,” which is not adequate to tackle complex diseases that involve multiple malfunctioned genes. The availability of big omics data offers new opportunities to transform the drug discovery process into a new paradigm of systems pharmacology that focuses on designing drugs to target molecular interaction networks instead of a single gene. Here, we develop a reliable multi-rank, multi-layered recommender system, ANTENNA, to mine large-scale chemical genomics and disease association data for the prediction of novel drug-gene-disease associations. ANTENNA integrates a novel tri-factorization based dual-regularized weighted and imputed One Class Collaborative Filtering (OCCF) algorithm, tREMAP, with a statistical framework that is based on Random Walk with Restart and can assess the reliability of a specific prediction. In the benchmark study, tREMAP clearly outperforms the single rank OCCF. We apply ANTENNA to a real-world problem: repurposing old drugs for new clinical indications that have yet had an effective treatment. We discover that FDA-approved drug diazoxide can inhibit multiple kinase genes whose malfunction is responsible for many diseases including cancer, and kill triple negative breast cancer (TNBC) cells effectively at a low concentration (IC_50_ = 0.87 μM). The TNBC is a deadly disease that currently does not have effective targeted therapies. Our finding demonstrates the power of big data analytics in drug discovery, and has a great potential toward developing a targeted therapy for the effective treatment of TNBC.

## 1 INTRODUCTION

The cost of bringing a drug to market has risen to approximately 2.6 billion dollars (Tufts Center for the Study of Drug Development, 2015), and the failure rate is daunting: only about one third of drugs in phase III clinical trials reach the market. The limited success of the conventional drug discovery process is largely attributed to the wide adoption of a reductionist model of “one-drug-one-gene-one-disease”[1-3]. As a matter of fact, the onset and progress of many complex diseases such as cancer is a systematic process that involves multiple interacting genes. Thus, it is necessary to design drugs that target gene interaction networks instead of a single gene. Moreover, drug repurposing that reuses existing safe drugs to treat new diseases has emerged as a new paradigm to accelerate drug discovery and development. As the safety profile of existing medicines has already been well documented, the cost of clinical trials can be significantly reduced.

Recent advances in high-throughput technologies have generated abundant chemical genomics data on drug actions and disease genes. These big, complex, heterogeneous data sets provide unprecedented opportunities for identifying genome-wide drug-gene-disease associations, thereby facilitating multi-targeted drug design and drug repurposing. However, several challenges remain in mining chemical genomics and disease association data for drug discovery. Firstly, chemical genomics data from high-throughput screening campaigns are not only extremely large but also highly noisy, biased, and incomplete. Many existing data mining algorithms cannot be directly applied to model chemical genomics data. Secondly, drug action is a complex process. It starts with drug-gene interactions at the molecular level, and manifest clinical outcomes through biological network. A single genomics data set can only capture one part of whole drug process. Thus, it is necessary to integrate multiple data sets for chemical-gene interactions, gene-disease associations, and chemical-disease associations to model the drug action on a multilayer. Finally, one of the fundamental problems in biomedical data mining has not been fully addressed: how to assess the individual reliability of a specific prediction from a data mining agent under a rigorous statistics framework. The reliable and unbiased assessment of the prediction quality for an individual instance is critical for cost-sensitive drug discovery process. For example, the selection of a novel chemical that is structurally different from patented drugs as a lead compound from a ranked list of candidate chemicals is a billion-dollar decision. Information on the individual predictive reliability of a novel chemical entity based on its weak chemical similarity to existing drugs in terms of bioactivity is invaluable. Most existing data mining tools can only provide an average predictive accuracy based on the population of training data, but not reliability for a specific new case. For example, in a ranking system, it is not straightforward to determine what the threshold is to select top-ranked hits. For a specific case, the top-first ranked hit could be a false positive. In another scenario, the top-*N* (*N*>1) ranked hits could all be true positives.

## 2 CONTRIBUTIONS OF THIS WORK

To address challenges in the predictive modeling of drug-gene-disease associations as well as unmet needs in the treatment of complex diseases such as cancer, this work makes contributions to both methodology development and translational medicine.

On the side of methodology development, our contribution is twofold. First, we have developed a novel algorithm tREMAP on the basis of tri-factorization to optimize matrix completion problem in which row and column have significantly different ranks. tREMAP formulates the chemical-gene predictions as a multi-rank dual-regularized weighted and imputed One Class Collaborative Filtering (OCCF) problem. Under the formulation of OCCF, negative data are not needed for the training, which are sparse and even unavailable. By using element-specific weights and imputation, tREMAP can handle noisy chemical genomics data in which the label is often uncertain. Finally, unlike conventional OCCF algorithm that applies a single rank to all layers, tREMAP assigns a different rank to a different layer. It is important since different layers can have dramatically different dimensions thus optimal ranks. For example, the dimension of a chemical layer is in the order of millions, while the dimension of a gene layer is only thousands. Our benchmark studies clearly show that tREMAP outperforms single-rank OCCF method. Second, to tailor the nature of chemical-gene-disease association data sets where observed chemical-disease associations are far sparser than known chemical-gene interactions and few three-way chemical-gene-disease associations exist, we have developed a multi-rank, multi-layered framework ANTENNA for inferring novel chemical-gene-disease associations. ANTENNA has three main components. (1) ANTENNA integrates multiple chemical genomics and disease association data set, and links them as a multi-layered network [4], as shown in Figure 1. (2) ANTENNA uses tREMAP to infer genome-wide novel chemical-gene associations. (3) Based on the genome-wide chemical-gene association, ANTENNA applies Random Walk with Restart (RWR) and a statistics framework, Enrichment of Topological Similarity (ENTS) [5], to predict chemical-disease associations and assess their reliabilities.

**Figure 1:**
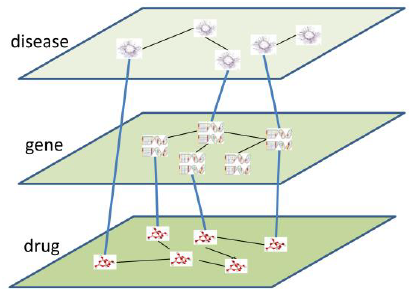
Illustration of Multi-Layered Network Model (MULAN) that integrates multiple genomics data sets.

Arguably, the most important contribution of this work is to discover a potentially safe and effective targeted therapy for triple negative breast cancer (TBNC). Using ANTENNA, we predicted that a FDA-approved drug diazoxide may inhibit multiple kinase genes. The malfunction of kinases is associated with many diseases such as cancer and Alzheimer’s disease. Among the kinases with the highest percentage of inhibition by diazoxide, one gene TTK is specifically over-expressed in the patients with TNBC [6, 7]. Thus, we hypothesized that diazoxide may kill TNBC cells. Our predictions were supported by multiple experimental evidences. TNBC is a subgroup of breast cancers, which is associated with the most aggressive clinical behavior. No targeted therapy is currently available for the treatment of TNBC. Our finding has a great potential toward developing a targeted therapy for the effective treatment of TNBC.

## 3 RELEVENT WORK

In principle, tensor factorization is a powerful method to infer three-way relationships. However, observed three-way chemical-gene-disease relations are extremely sparse. Majority of observed chemical-gene pairs are not associated with any diseases. Thus, the tensor factorization may be not the best option for this work. OCCF has been applied to a bipartite graph for predicting drug-target interactions[8], but not to inferring multiple drug-gene-disease associations. Moreover, existing OCCF algorithm is mainly based on the formulization of matrix factorization that only allows a common rank for both row and column. FASCINATE is an algorithm that can jointly infer missing links from a multi-layered network model[4]. However, FASCINATE is based on the formulation of a single rank collective OCCF. Moreover, it can only rank predicted relations[4]. There is no reliability information associated with each individual prediction. This work will address the drawbacks in matrix factorization, OCCF, and FASCINATE when applied to inferring chemical-gene-disease associations.

## 4 EXPERIMENTAL AND COMPUTATIONAL DETAILS

### 4.1 Overview of Computational and Experimental Procedure

Our primary purpose is to mine chemical genomics and disease association data to identify novel targeted therapies for unmet biomedical problems such as the treatment of TBNC. As shown in Figure 2, the input of ANTENNA is the existing chemical genomics, drug, and disease databases including DrugBank[9], ZINC[10], ChEMBL[11], and CTD[12]. We first integrate these data sets into a multi-layered chemical-gene-disease network, MULAN. Then we apply tREMAP, a multi-rank dual-regularized weighted imputed OCCF algorithm, to infer novel chemical-gene associations. Next, we used ENTS to predict drug-disease association and to assess the reliability for each inferred association. The output of ANTENNA is a list of ranked drug-disease associations ranked by their statistical significance. Finally, we experimentally validate the top-ranked predictions.

**Figure 2:**
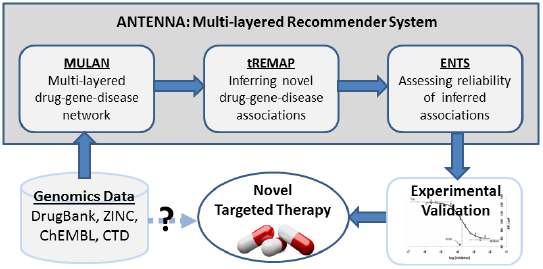
Workflow of drug discovery process using ANTENNA, a multi-layered recommender system.

### 4.2 Construction of Multi-Layered Chemical-Gene-Disease Network (MULAN)

We integrated heterogeneous data sets from genomics into a multi-layered network model, MULAN. In the MULAN, each node is a chemical entity (drugs and other chemicals), a biological entity (genes or proteins that it encodes), or a phenotypic entity (disease and side effect). Nodes in the same entity class are linked together by similarities (e.g., chemical-chemical similarity) or interactions (e.g., protein-protein interactions). Nodes that belong to different entity classes reside in different network layers and are linked by known associations (e.g., drug-target interactions, disease-gene associations). Integration of genomics data into a bipartite graph is of proven value[13]. The MULAN can be considered as the unification of multiple bipartite graphs; thus, our new method is likely to be more robust than traditional approaches.

Chemical-gene associations including drug-gene associations were obtained from the ZINC [14], ChEMBL [15] and DrugBank [16] databases. To obtain reliable chemical-gene association pairs, binding assays records with IC_50_ information were extracted from the databases, and the cutoff IC_50_ value of 10 µM was used where applicable. Chemical-gene pairs were considered associated if IC_50_≤10 µM (active pairs), unassociated if IC_50_>10 µM (inactive pairs), ambiguous if records exist in both ranges (ambiguous pairs), and unobserved otherwise (unknown pairs). A total of 198,712 unique chemicals and 3,549 unique genes were obtained from the combination of ChEMBL and ZINC with 228,725 unique chemical-gene active pairs, 76,643 inactive pairs, and 4,068 ambiguous pairs. Of the 198,712 chemicals, 722 were found to be FDA-approved drugs. Furthermore, drug-gene relationships were extracted from the DrugBank and integrated into the ZINC_ChEMBL dataset above. A total of 199,338 unique chemicals and 6,277 unique genes were obtained from the combination of ZINC, ChEMBL, and DrugBank with 233,378 unique chemical-gene active pairs. Drug-disease and gene-disease associations were directly obtained from the Comparative Toxicology Database (CTD) [12].

Chemical-chemical similarity scores are one of the required inputs of tREMAP. Although there are a number of metrics developed for chemical-chemical similarity, a recent study showed that Jaccard index-based similarity is highly efficient for fingerprint-based similarity measurement [17]. The fingerprint of choice in this study is the Extended Connectivity Fingerprint (ECFP), which has been successfully applied to chemical structure-based target prediction method, PRW [18]. Jaccard index is used to calculate a similarity score between two chemicals, c_1_ and c_2_.

Gene-gene similarity scores are also one of the required inputs for tREMAP. The similarity between two proteins encoded by genes was calculated based on their amino acid sequence similarity using NCBI BLAST [19] with an e-value threshold of 1×10^−5^ and its default options. A similarity score for query protein *p*_1_ to target protein *p*_2_, *d*_*bit* (*p*1,*p*2)_, was calculated by the ratio of a bit score for the pair compared to the bit score of a self-query. To be specific, for the query protein *p*_1_ to the target protein *p*_2_, protein-protein similarity score was defined such that *T*_(*p*1,*p*2)_ = *d*_*bit* (*p*1,*p*2)_*/d*_*bit* (*p*1,*p*1)_.

Disease-disease similarity is required for tREMAP to infer chemical-disease associations, and can be calculated using distributed word representation[20]. In this work, we do not infer the chemical-disease association directly using tREMAP, since only less than 0.4% of chemicals have observed associations with one or more diseases. Instead, we use ENTS and target binding profile of a chemical, which is derived from tREMAP, to infer the chemical-disease associations.

### 4.3 tREMAP Algorithm

Our prediction method tREMAP is based on a tri-factorization one-class collaborative filtering algorithm. In the case of chemical-gene association, it assumes that similar chemicals will interact with similar genes, and unobserved associations are not necessarily negative. Assuming that a fairly low number of factors (i.e. smaller number of features than the number of total chemicals or genes) may capture the characteristics determining the drug-gene associations, two low-rank matrices, *F* (drug side) and *G* (gene side), were approximated such that 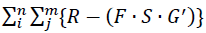 is minimized where *R* is the matrix for known drug-gene interactions and *G*′ is the transposition of the gene side low-rank matrix *G*. The two low rank matrices, *F*_*n*×*r*1_ with the rank of *r1* and *G*_*m*×*r*2_ with the rank of *r2*, and their connectivity matrix *S*_*r*1×*r*2_ are obtained by iteratively minimizing the objective function,

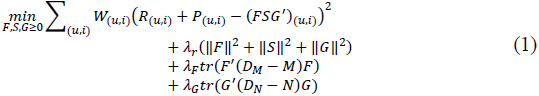

Here, *W*_(*u,i*)_ is the penalty weight on the observed and unobserved associations which indicate the reliability of the assigned probability of true association, *P*_(*u,i*)_ is the imputed value (i.e. the probability of unobserved associations as real associations), *M* and *N* is the symmetric chemical-chemical similarity matrix and gene-gene similarity matrix, respectively. *D*_*M*_ and *D*_*N*_ are the degree matrix of *M* and *N*, respectively. λ_*r*_ is the regularization parameter to prevent overfitting, λ_*F*_ is the importance parameter for chemical-chemical similarity, λ_*G*_ is the importance parameter for gene-gene similarity, and *tr*(A) is the trace of matrix A. The weight and imputation values can be determined by *a priori* knowledge or from the prediction of other machine learning algorithms.

The optimization problem defined in Eq. (1) is non-convex. Thus, we seek to find a local optimum by the block coordinate descent method. In Eq. (1), *D*_*M*_, *M, D*_*N*_, and *N* are non-negative matrices. The derivative Eq. (1) with regard to F, G, and S with the non-negativity constraint has a fixed-point solution. To scale up tREMAP in terms of both time and storage, we propose efficient multiplicative updating rules as follows:

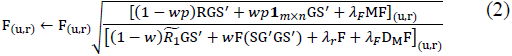

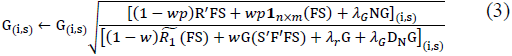

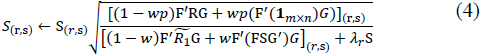

Where *w* and *p* are weighted and imputed value, respectively. They are either set based on a priori knowledge (e.g. the false positive rate of a high-throughput screening experiments), or can be tuned as hyper-parameters. 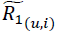 is the sparse matrix in which the value of elements is predicted by *F* and *G* on the observed cases Θ in *R*, i.e.

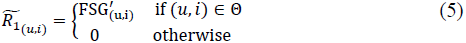

We use a block-coordinate descent algorithm to iteratively update F, G, and S.

The raw predicted score for the *i*th chemical to bind the *j*th protein can be calculated by *P*_(*i,j*)_ = *F*_(*i*,:)_ · *S* · *G*_(*j*,:)_′. Also, the matrix *F*_*n*×*r*1_ is referred to as a low-rank drug profile since its *i*th row represents the *i*th drug’s behavior in the drug-gene association network as well as drug-drug similarity spaces compressed to *r*1 number of features.

### 4.4 ENTS Algorithm

#### 4.4.1 Overview of ENTS algorithm

The rationale of ENTS is that when clusters of instance share common features, a cluster ranked closely together is more likely similar to the new instance than a cluster ranked randomly or spread out across the ranking. In addition, network topological similarity provides more robust and accurate global ranking across an entire hypothesis space than pairwise similarity does. Unlike conventional local ranking (e.g., *k*-nearest neighbors), global instance ranking can support statistical enrichment analysis because it draws valuable information on the ranking for all instances in a cluster from lower, non-randomly ranked cases.

#### 4.4.2 Classification or clustering of database instances

To initialize ENTS, part or all of the instances in the database (training set) are classified based on target feature *T*. In ANTENNA, the *T* is the disease associated with a drug. If database instances are not pre-classified, clusters of training data are assembled using *T* features under unsupervised clustering techniques [21] such as *k*-means [22], mean-shift [23], affinity propagation [24], or *p*-median model [25] etc. After the classification or clustering, each instance cluster will be assigned with a unique label (i.e. a specific disease in ANTENNA). These instance clusters are applied to the next step. It is noted that the instance clusters are not necessarily disjointed. They can overlap.

#### 4.4.3 A weighted graph represents training instance similarity by T-features

After the initialization, ENTS builds a database instance graph; a weighted graph with one node for the *T*-feature of each training instance and an edge between two nodes only if their pairwise similarity exceeds a certain threshold. The threshold depends on the features and the pairwise similarity metric. Any similarity metric (e.g. Euclidean distance, Jaccard index, Hidden Markov Model, kernel-based similarity etc.) can be applied here. In ANTENNA, we use cosine similarity of low-rank profile of drugs to measure the distance between drugs.

#### 4.4.4 Network topological similarity

Given a query with known *K*-feature and the goal to predict its unknown *T*-feature, ENTS first links the query to all nodes in the training instance graph, where new edges are not found in the training instance graph. The weights of these new edges are only based on *K*-feature similarity. Then Random Walk with Restart (RWR) is applied to perform a probabilistic traversal of the instance graph across all paths leading away from the query, where the probability of choosing an edge will be proportional to its weight. The algorithm will output a list of all instances in the graph, ranked by the probability *tiq* that a path from the query will reach the node *i*. In this way, RWR can capture global relationships that may be missed by pairwise similarity [26].

We modified the RankProp algorithm [27], a variant of RWR. The graph is represented as an adjacency list to save memory and speed up the iterative algorithm. The current implementation is scalable to a graph with millions of nodes and hundreds of millions of edges.

#### 4.4.5 Statistical significance of network topological similarity

A network topological search only ranks instances based on their similarity, but gives no information on the reliability of the ranking. To assess the statistical significance of the ranking of an instance cluster *C*_*i*_ generated previously, ENTS compares the score distribution of the cluster *C*_*i*_ with that of a randomly drawn cluster of the same size. When the mean of global topological similarity scores 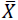 in a cluster is used as the statistic, an efficient random-set method is used for the parametric approximation of the null distribution [28]. The random-set method compares an enriched cluster of size *m* with all other distinct clusters of size *m* drawn randomly from a case graph on *N* nodes. The exact distribution of 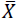 is intractable, but can be approximated with the normal distribution with mean and variance as follows:

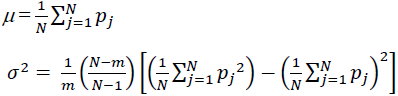

Where *p*_*j*_ is the global topological similarity score of the structure *j* in the graph to the query. The enrichment score of the cluster *C*_*i*_ is then normalized with

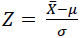

A p-value and Benjamini-Hochber adjusted false discovery rate (FDR) is then calculated for each *Z*-score.

### 4.5 Combining tREMAP and ENTS to Predict Drug-Disease Association

In ANTENNA, we firstly use tREMAP to generate chemical-side low rank matrix *U*_*UP*_ and gene side low-rank matrix *VUP*. The *i*th row of *U*_*UP*_ contains the gene association profile for the *i*th drug. Then, we calculated drug-drug cosine similarities based on the matrix *UUP*, and construct a drug-drug similarity graph. For each row of *U*_*UP*_ for FDA approved drugs, the cosine similarity of drug c1 and drug c2 can be calculated by,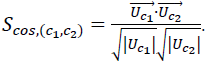 To search for possibly undiscovered uses of the drugs, we focus on drugs that are found to have high cosine similarity but low chemical structural similarity (< 0.5). Finally, we cluster drugs based on their directly or indirectly associated diseases annotated in CTD database[12], and use ENT to assess and rank the statistical significance of novel drug-disease associations. The final output of ANTENNA is the ranked list of predicted drug-disease association based on FDR.

### 4.6 Experimental Validation

#### 4.6.1 Kinase binding assay

Kinase is an enzyme that catalyzes the transfer of a chemical group phosphate to another biomolecule. It functions as a molecular switch in many biological processes. The malfunction of kinases is responsible for many diseases such as cancer. There are more than 400 kinases in the human genome, which is termed as kinome. To rigorously validate the performance of ANTENNA, we employed a competition binding assay to detect the binding of selected drugs to a set of 438 kinases (human kinome). The proprietary KinomeScan^TM^ assay was performed by DiscoverX (CA). The assay tested the capacity for a drug to disrupt the binding of each DNA-tagged kinase to a support which one was in turn bound to the kinase’s known ligand. If binding between the kinase and its known ligand was disrupted in the presence of the drug, this indicated that the drug either competed directly with the known ligand or allosterically altered the kinase’s ability to bind to that ligand. DMSO was used as a positive control and a pico-molar kinase inhibitor was used as a negative control. Binding levels were quantitated by performing real-time polymerase chain reaction (qPCR) on the DNA tag of the ligand-bound kinases. The qPCR is a molecular biology technique to amplify a single copy or a few copies of DNA segment in several orders of magnitude, and to measure the reaction in a real time. The tests were performed at 100 μM concentration of tested drug, and results were reported as %Control, calculated as follows:

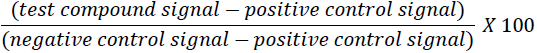

A lower %Control score indicates a stronger interaction.

#### 4.6.2 Cancer cell viability assay

MCF-7 cells from ATCC^®^ and MDA-MB 468 cells (a gift of Dr. R Sullivan from Queens Community College, the City University of New York) were used for this study. MCF-7 is breast cancer cell line. MDA-MB 468 is triple negative breast cancer cell line which does not express estrogen receptor (ER), progesterone receptor (PR), and human epidermal growth factor receptor (Her2/neu). Cells were cultured in Dulbecco’s Modified Eagle Medium (DMEM) (Thermo Fisher Scientific) supplemented with 10% fetal bovine serum (Thermo Fisher Scientific) and 50 μg/ml gentamicin (Thermo Fisher Scientific) at 37°C 5% CO_2_ incubator.

Cell viability was determined by neutral red assay which is based on the lysosome uptake of neutral red dye [29]. Briefly, cells (2 x 10^4^ cells per well) were plated onto 96-well plate in a total volume of 200 μl on the day before chemical treatments. Chemicals were dissolved in dimethyl sulfoxide (DMSO) to obtain 0.1 M stock solution 15 minutes before chemical treatments. Then various concentrations (0.1-150 μM) of chemicals were prepared in fresh media. The final concentration of DMSO in each well was equal to or less than 0.15% which is considered non-toxic to cells [30].

After 24 hours of chemical treatments, 20 μl of 0.33% Neutral Red Solution (Sigma Aldrich) was added onto wells. After 2 hours incubation at 37°C 5%CO_2_ incubator, dye solution was carefully removed and cells were rinsed with 200 μl Neutral Red Assay Fixative (0.1% CaCl_2_ in 0.5% formaldehyde) (Sigma Aldrich) twice. The absorbed dye was then solubilized in 200 μl of Neutral Red Assay Solubilization Solution (1% acetic acid in 50% ethanol) (Sigma Aldrich) for 10 minutes at room temperature on a shaker. Absorbance at 540 nm and 690 nm (background) was measured by BioTek Synergy Mx microplate reader.

Each concentration in each experiment was done with at least triplicate. Multiple experiments were done to obtain IC_50_ values for each drug and each cell line. The viability was determined based on a comparison with untreated cells which were set as 100% cell viability. The IC_50_ values which represent the chemical concentration needed to inhibit 50% cell proliferation were calculated from the dose-response curve.

## 5 RESULTS AND DISCUSSION

### 5.1 Performance evaluation of tREMAP

In our published studies[8], single rank REMAP outperformed state-of-the-art methods: a chemical similarity-based method (PRW [18]), the best performed matrix factorization methods so far (NRLMF [31] and KBMF with twin kernels (KBMF2K) [32]), combination of WNN and GIP (WNNGIP [33]), and another type of collaborative filtering algorithm (Collaborative Matrix Factorization (CMF) [34]). Here we compare the performance of tREMAP with that of REMAP using two benchmarks. The first benchmark includes 3,494 chemicals, 25 G-protein coupled receptors (GPCRs), and 4,494 observed chemical-GPCR associations. The second benchmark includes 33,684 chemical, 31 Cytochrome P4_50_ enzymes (CYP4_50_), and 51,699 observed chemical-CYP450 associations.

As shown in Figure 3, tREMAP clearly outperforms REMAP when evaluated by both of the benchmarks. tREMAP identifies around 96% and 87% true associations ranked on the top 3 for GPCR and CYP450, respectively, while REMAP can only identify around 78% and 60% true hits ranked on top 3 respectively.

**Figure 3:**
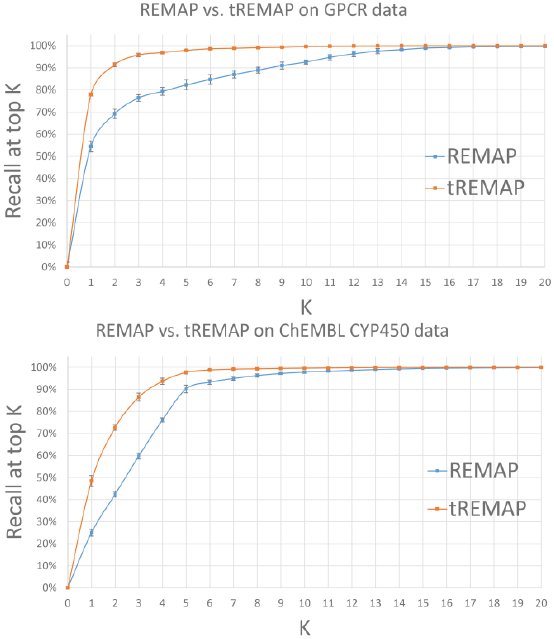
Performance comparison of tREMAP with REMAP for GPCR (upper) and CYP450 (bottom), respectively. Performance is measured by the recall at the top rank *K*.

When evaluated by the application to sequence-structure similarity search, ENTS is superior to Hidden Markov Model and RWR[5].

### 5.2 Time complexity of tREMAP

Empirically, the running time of tREMAP is linearly dependent on the number of chemicals and genes, as shown in Figure 4. When evaluated in a machine with 2 cores of 2.18 GHz CPU. It takes around 1,000 seconds for a matrix with 15,000 chemicals, 200 genes, chemical-side rank of 1,000, and gene-side rank of 200 to converge.

**Figure 4:**
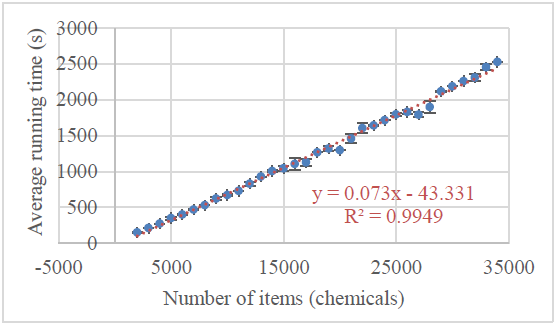
Running time of tREMAP vs the number of items. The computational time was measured in a 2 cores of 2.18 GHz CPU, for a matrix with 200 genes and varied number of chemicals. The ranks for chemical and gene are fixed as 1,000 and 200, respectively.

### 5.3 ANTENNA Predictions

By combining tREMAP with ENTS, ANTENNA predicted that 21,921 novel drug-disease associations with Benjamini-Hochberg adjusted false discovery rate (FDR) less than 0.02. We selected a drug-disease pair for further experimental evaluation based on the following criteria. First, the drug was predicted to bind kinases, as the genome-wide binding assay for kinases is accessible. Second, the associated disease does not have effective therapy, so that the repurposed drug will have the biggest clinical impact. Third, the cell-based disease model is available, so that we can evaluate the efficacy of the drug.

Based on above criteria, diazoxide, a safe FDA-approved drug for hypertension, was selected. Diazoxide was predicted to interact with protein kinases. Furthermore, ANTENNA predicted that diazoxide was associated with Triple Negative Breast Cancer (TNBC) with Benjamini-Hochber adjusted false discovery rate (FDR) of 0.0108. Thus, diazoxide may be repurposed for the treatment of TBNC which is the most aggressive type of breast cancer and cannot be treated by any existing targeted therapy. It notes that the FDR of predicted diazoxide-TNBC association is not particular statistically significant. If this prediction is experimentally validated, we will have more confidence in predictions with lower FDRs.

### 5.4 Kinase Binding Assay

We validated the binding of diazoxide to kinases using KinomeScan^TM^ assay. Figure 5 displays the binding profile of diazoxide across 438 kinases (kinome). Diazoxide has the highest percentage inhibition of kinases DRYK1A, IRAK1, and TTK with 7.0%, 8.9%, and 15.0% control. It is noted that the lower %Control, the higher inhibition of kinase activity.

**Figure 5:**
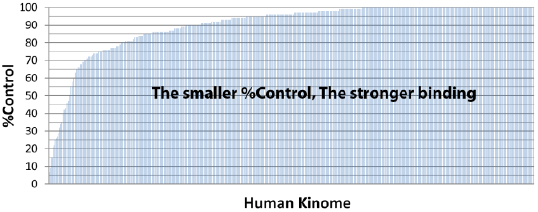
Binding profile of FDA-approved drug diazoxide (100μM) on 438 kinases determined by KinomeScanTM assay.

As shown in Table 1, the malfunction of DYRK1A, IRAK1, and TTK is associated with multiple diseases, especially cancers and Alzheimer’s disease. To verify our predictions, we tested the effect of diazoxide on breast cancer cells.

**Table 1:** Gene-disease associations of three kinases that have the highest inhibition percentage by diazoxide

| Kinase | KinomeScan™ %Control | Gene-Disease Association |
| --- | --- | --- |
| DYRK1A | 7.0 | Multiple cancer drug-resistance, Alzheimer's disease |
| IRAK1 | 8.9 | Breast cancer metastasis, herpesvirus lymphoma, Alzheimer's disease |
| TTK | 15 | TNBC, Hepatocellular Carcinoma |

### 5.5 Cancer cell viability assay

The cytotoxicity of diazoxide was determined by neutral red cell viability assay. The IC_50_ values obtained from Estrogen positive breast cancer MCF-7 cells and TNBC MDA-MB-468 cells treated with chemicals for 24 hours were shown in Table 2. Diazoxide was much more effective in inhibiting the cell proliferation of TNBC cancer MDA-MB 468 cells as compared to MCF-7 breast cancer cells with the values of IC_50_ 0.87 ± 0.39 ±M and 130.0 ± 70.0 ±M, respectively. The IC_50_ is the concentration of diazoxide that inhibits the cell proliferation of 50% cancer cells. The smaller the IC_50_ value is, the stronger anti-cancer activity diazoxide has. It is accepted that a chemical compound is active when the IC_50_ is less than 10 ±M. Thus, diazoxide could be a highly effective targeted therapy for the treatment of TNBC at a low concentration.

**Table 2:**
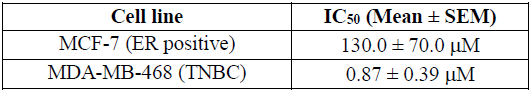
IC_50_ values of diazoxide on cancer cells

## 6 CONCLUSIONS

In summary, we have developed a reliable and accurate multi-rank, multi-layered recommender system ANTENNA. Using ANTENNA, we predicted that FDA-approved safe medicine diazoxide could bind to kinases whose malfunction is associated with TNBC. KinomeScanTM assay confirmed the kinase binding of diazoxide. Cancer cell viability assay further validated that diazoxide is highly effective in inhibiting the proliferation of TNBC cancer cells. These findings suggest that diazoxide can be repurposed as an effective targeted therapy for the treatment of TNBC. Furthermore, diazoxide may be effective in the treatment of other diseases such as hepatocellular carcinoma and Alzheimer’s disease. We are carrying out experiments to verify these predictions. This study demonstrates that big data analytics provides new opportunities for accelerating drug discovery and development, and realizing the full potential of precision medicines.

## ACKNOWLEDGMENTS

This work was partly supported by Grant Number R01LM011986 from the National Library of Medicine (NLM) of the National Institute of Health (NIH), Grant Number R21TR001722 from the National Center for Advancing Translational Sciences of NIH, and Grant Number MD007599 from the National Institute on Minority Health and Health Disparities (NIMHD) of NIH.

